# Same Equilibrium. Different Kinetics. Protein Functional Consequences

**DOI:** 10.1101/838938

**Authors:** Sonja Schmid, Thorsten Hugel

**Affiliations:** Institute of Physical Chemistry, University of Freiburg, Germany; Department of Bionanoscience, Kavli Institute of Nanoscience Delft, Delft University of Technology, The Netherlands; Signalling research centers BIOSS and CIBSS, Albert Ludwigs University, Freiburg, Germany

## Abstract

In a living cell, protein function is regulated in several ways, including post-translational modifications (PTMs), protein-protein interaction, or by the global environment (e.g. crowding or phase separation). While site-specific PTMs act very locally on the protein, specific protein interactions typically affect larger (sub-)domains, and global changes affect the whole protein in non-specific ways.

Herein, we directly observe protein regulation in three different degrees of localization, and present the effects on the Hsp90 chaperone system at the levels of conformational equilibria, kinetics and protein function. Interestingly using single-molecule FRET, we find that similar functional and conformational steady-states are caused by completely different underlying kinetics. Solving the complete kinetic rate model allows us to disentangle specific and non-specific effects controlling Hsp90’s ATPase function, which has remained a puzzle up to this day. Lastly, we introduce a new mechanistic concept: functional stimulation through conformational confinement. Our results highlight how cellular protein regulation works by fine-tuning the conformational state space of proteins.

**Significance:** Proteins are perceived more and more as dynamic systems whose function depends critically on local and global flexibility. While 3D structures of proteins are frequently available today, our models often lack the time component, namely rate constants that determine protein function and regulation.

Here we used single-molecule FRET to elucidate how the chaperone protein Hsp90 is regulated on various levels, locally and globally. We find that ATPase stimulation occurs not only through specific interactions, but also non-specifically by reducing non-productive conformational flexibility; i.e. by changing kinetics rather than thermodynamics. Our work introduces ‘stimulation through conformational confinement’ as a general mechanistic concept. We anticipate that this concept plays an important role in protein regulation, phase separation, and in dynamic protein systems in general.

## Maintext

Protein function is essential for life as we know it. It is largely encoded in a protein’s amino-acid chain that dictates not only the specific 3D structure, but also the conformational flexibility and dynamics of a protein in a given environment. Precise regulation of protein function is vital for every living cell to cope with an ever-changing environment, and occurs on many levels pre- and post-translationally (1,2). After translation by the ribosome, protein function depends strongly on post-translational modifications (PTMs) (3), but also on binding of nucleotides (4), cofactors (5), various protein-protein interactions (PPIs) (6), and global effects, such as temperature (7), macro-molecular crowding and phase separation (8), redox conditions (9), osmolarity (10) etc. Importantly, this regulation occurs on very diverse levels of localization. Global effects affect the whole protein non-specifically, PPIs act at a given interface and site-specific modifications are very localized. Nevertheless, all of them influence the molecular properties that determine the 3D conformation, the conformational dynamics, and thereby also the function of a protein (11–15). The chaperone protein Hsp90 (16) is an excellent test system to investigate diverse regulation mechanisms (17). It was recently discussed that a single PTM can functionally mimic a specific co-chaperone interaction in human Hsp90 (18). Here we take a next step and disentangle how a PTM-related point mutation, a co-chaperone interaction, and macro-molecular crowding affect the function, kinetics, and thermodynamics of this multi-domain protein.

Hsp90 is an important metabolic hub. Assisted by about twenty known cochaperones, yeast Hsp90 is involved in the maturation of 20% of the entire proteome (19). Among its substrates (referred to as ‘clients’) are many kinases involved in signal transduction, hormone receptors, the guardian of the genome p53 (20), but also cytoskeletal proteins, e.g. actin, tubulin, and many more (21,22). Cancer cells were found to be ‘addicted’ to Hsp90 (23), which is therefor also a prominent drug target in cancer research. Hsp90 is a homo-dimer where each monomer consists of three domains (24): the N-terminal domain (N) with a slow ATPase function, the middle domain (M) believed to be the primary client interaction site (25), and the C-terminal domain providing the main dimerization contacts. Apart from closed conformations, where the three domains align in parallel, Hsp90 exists primarily in v-shaped, open conformations with dissociated N-terminal and middle domains (26,27). Both global arrangements are semi-stable at room temperature. As a consequence, Hsp90 alternates constantly between open and closed conformations - even in the absence of the chemical energy source, ATP (28–30). Surprisingly, the characteristic conformational changes of Hsp90 are only little affected by e.g. anti-cancer drug candidates (31) or natural nucleotides (29). In addition, to the stress-induced isoform discussed herein (Hsp82), there is also a cognate isoform (Hsc82) in yeast, which differs in unfolding stability, client range etc. despite 97% sequence identity (32).

Here we present three orthogonal ways to modulate Hsp90’s conformational state space, illustrated in **Fig. 1**. The investigated point mutation A577I is located in the C-terminal hinge region of Hsp90. Residue A577 is the equivalent of a post-translational S-nitrosylation site in human Hsp90 (33). While nitrosylation of that residue has a two-fold inhibitory effect – on the ATPase function and the client stimulation by human Hsp90 – the A577I mutation caused a nearly 4-fold amplification of the ATPase rate (34). The fact that the point mutation is located far away from the ATP binding site indicates a long-range communication from the C-domain all the way to the N-terminal ATPase site, offering valuable, mechanistic insight in Hsp90’s intra-molecular plasticity. Second, we consider the protein-protein interactions between Hsp90 and the activating co-chaperone Aha1, which is a well-known stimulator of Hsp90’s inherently slow ATPase activity. It makes contacts to the middle and N-terminal domain, which rearranges the ATP lid (35), and the catalytic loop (including Arg380) (36) in a favorable way for ATP hydrolysis. The affinity of Aha1 for Hsp90 itself is also markedly enhanced by PTMs (37). The third way of modulation mimics the crowding encountered in the cell, which is full of proteins, nucleic acids, vesicles and organelles. We mimic cellular macro-molecular crowding using the common crowding agent Ficoll400, i.e. branched polymeric sucrose. In contrast to the previous two modulations, crowding represents a completely nonspecific, physical interference.

**Fig. 1:**
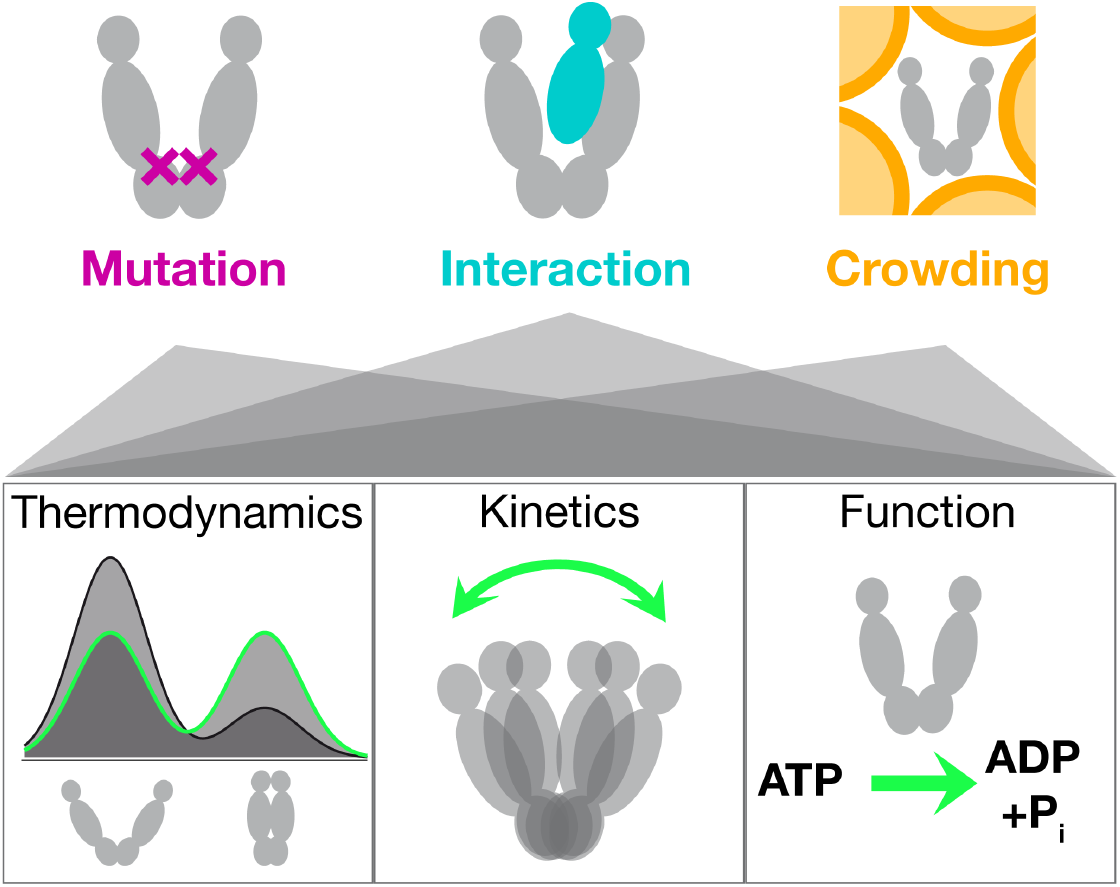
Protein regulation uses different degrees of localization. Mutations or PTMs act most locally, protein-protein interactions (PPIs) act on the protein domain level, and changes in the global environment, such as crowding or phase separation, act non-specifically and globally on the protein. Each of them affect conformational thermodynamics and kinetics to fine-tune the protein conformational state space and thereby protein function.

At first sight, all three modulations provoke a similar steady-state behavior in Hsp90. But our single-molecule experiments allow us to disentangle the different underlying causes thereof.

## Results

### Mutation, cochaperone and crowding show similar thermodynamics

First we follow yeast Hsp90’s conformational kinetics in real-time using single-molecule Förster resonance energy transfer (smFRET) measured on a total-internal reflection fluorescence (TIRF) microscope (**Fig. 2a**). The FRET pair configuration displayed in **Fig. 2a** (top) results in low FRET efficiency (little acceptor fluorescence) for v-shaped, open conformations of Hsp90, and high FRET efficiency (intense acceptor fluorescence) for more compact, closed conformations. This allows us to obtain steady-state information, like the population of closed conformation, and also the kinetics of conformational changes (29,38–40). Example traces obtained from three Hsp90 molecules under different conditions are displayed in **Fig. 2b**: for the point mutant A577I, in the presence of the cochaperon Aha1, or under macro-molecular crowding by Ficoll400. The observed transitions between the low- and high-FRET states reflect global opening or closing. **Fig. 2c** shows the steadystate population of open and closed conformations. In all three cases, a shift towards closed conformations is observed with respect to the corresponding reference distribution obtained under equivalent experimental conditions (see Methods for details).

**Fig. 2:**
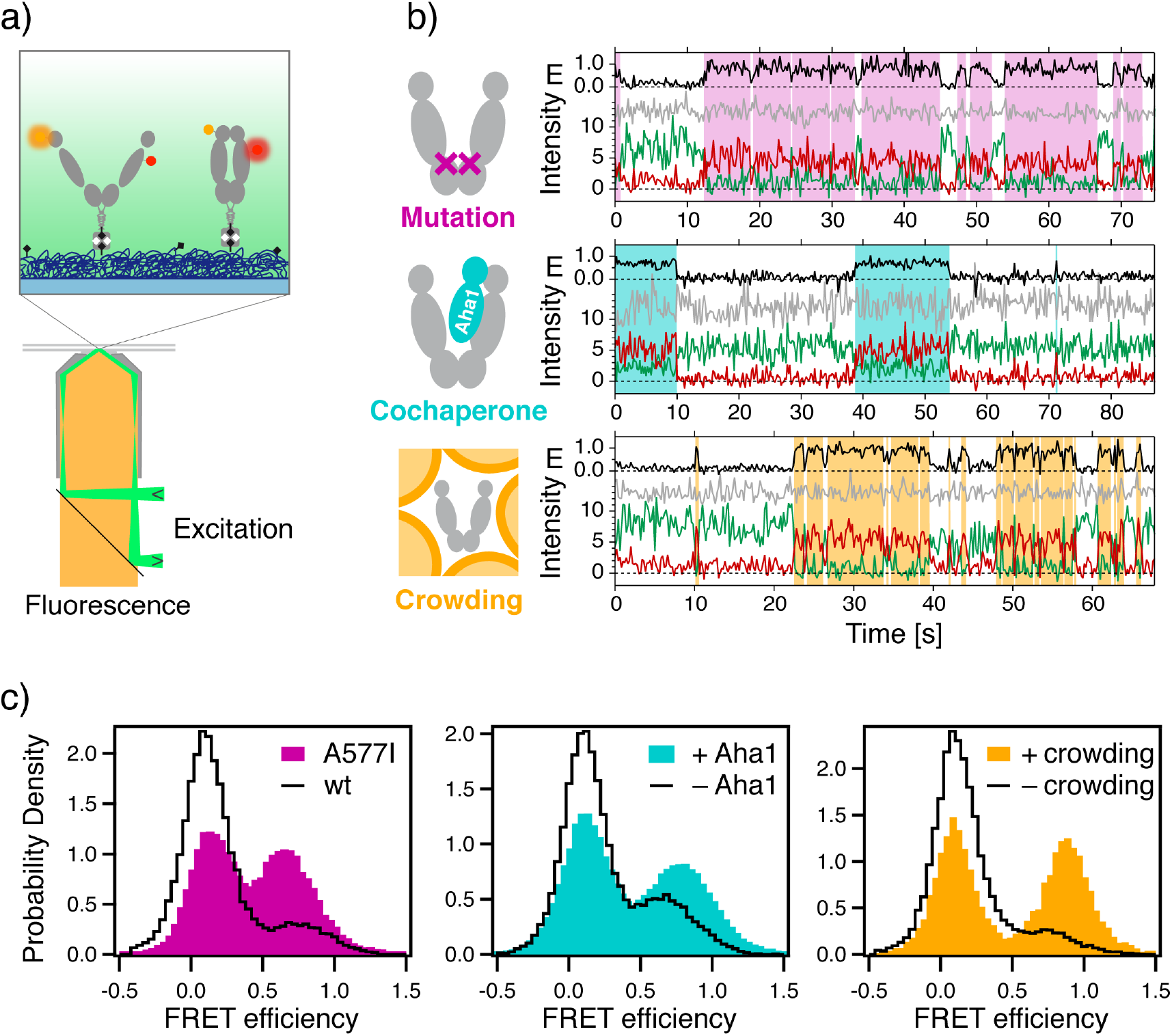
Mutation, cochaperone interaction, and crowding show similar thermodynamic effects. (a) Illustration of the single-molecule FRET experiment using an objective-type TIRF microscope (bottom): cross section through the objective, and flow chamber (both gray) and the dichroic mirror (black) separating the laser excitation (green) from the collected fluorescence (yellow). The zoom view (top) shows the fluorescently labeled Hsp90 (FRET donor, orange; acceptor red), which is immobilized on a PEG-passivated (dark blue) coverslip (light blue) using biotin-neutravidin coupling (black and gray). (b) Example time traces obtained from individual Hsp90 molecules for the point mutant A577I (top), in the presence of 3.5μM cochaperone Aha1 (center), or under macro-molecular crowding by 20wt% Ficoll400: FRET efficiency E (black), fluorescence of the FRET donor (green), acceptor (red), directly excited acceptor (gray). White and colored overlays denote low- and high-FRET dwells, respectively, as obtained using a hidden Markov model and the Viterbi algorithm. (c) FRET histograms compiled from many singlemolecule trajectories as indicated, and normalized to unity (wt: wild type). Reference data (black) was measured under the specific conditions of each of the three experiment series (see Methods). The number of individual molecules included per histogram are: A577I, 181; wt, 163; +Aha1, 122; −Aha1, 231; +crowding, 50; -crowding, 81.

The A577I mutation increased the closed population from 16% to 47%. This is a large change considering that this hydrophobic-to-hydrophobic mutation is not a drastic change to the overall charge distribution. In particular, as the slightly bulkier isoleucine side chain points outward in the crystal structure of Hsp90’s closed conformation (24). In addition to the A577I homodimer, already the A577I/wild type (wt) hetero-dimer shows a considerably larger population of closed conformations, especially in the presence of ATP (**Fig. S1** left). Under ADP conditions the additive effect is also observed, but weaker (**Fig. S1** right). Under both conditions, the second A577I in the homodimer leads to a further shift towards closed conformations. The slight but consistent shift of the corresponding low-FRET peak in **Fig. 2c** (left) could be explained by a sterical hindrance of the farthest opening in the A577I homodimer. Furthermore, very fast transitions at the temporal resolution limit (200ms) occurred more frequently, which can be seen by the increased overlap between the two populations. Both, sterical hindrance and faster transitions, can be interpreted as a global stiffening of Hsp90’s structural core formed by the C- and middle domain. The interaction with Aha1 (**Fig. 2c**, center) forms inter-domain (N-M) and inter-monomer contacts. The latter increase the affinity for N-N binding, and thus cause Hsp90 to shift from 29% closed to 46% closed population, which is in line with previous qualitative reports (41,42). Lastly, macro-molecular crowding increased the closed population from 14% to 52%, in agreement with previous ensemble findings (43). In contrast, to the A577I mutant, crowding slowed down fast fluctuation at the resolution limit, which creates well-separated populations in **Fig. 2c**). The induced shift towards closed conformations appears in a concentration-dependent manner, as can be seen in **Fig. S2**. In contrast to polymeric, branched sucrose (Ficoll400), monomeric sucrose had only negligible effect on the population distribution. This proves that macro-molecular crowding is the cause of the observed population shift, and a biochemical glucose-associated reason can be dismissed.

In all three cases, a clear shift towards closed conformations is observed, although to slightly different extents. Importantly, based on these distributions alone, the *energetic* origin of the population shift remains unclear. I.e. whether the observations arise from a stabilization of closed conformations, or a destabilization of the open conformations, or even a combination of both. To answer these questions, we solved the full kinetic rate model, presented below.

### Same thermodynamics, different conformational kinetics

**Fig. 3a** shows the kinetic rate models describing Hsp90’s global opening and closing dynamics. The significant changes caused by each type of modulation are highlighted in red and green as indicated. The corresponding quantitative rate changes and confidence intervals are displayed in **Fig. 3b**. We used the Single-Molecule Analysis of Complex Kinetic Sequences (SMACKS (29)) to quantify rate constants and uncertainties directly from the smFRET raw data. For Hsp90’s global conformational changes, we consistently infer 4-state models (29,31,44): two low-FRET states (open conformations) and two high-FRET states (closed conformations). Although only two different FRET efficiencies can be resolved, at least four kinetic states are needed to describe the observed kinetic heterogeneity. Based on recent results (27), we expect an entire ensemble of open sub-conformations that - on the timescale of the experiment - are sufficiently well described by two kinetically different low-FRET states. The two closed states, one short-lived (state 2) and one longer-lived (state 3), likely differ in local conformational elements. The well-known N-terminal beta-sheet with or without its cross-monomer contacts (as observed in the closed crystal structure (24)) could explain the additional stabilization of state 3 with respect to state 2.

**Fig. 3:**
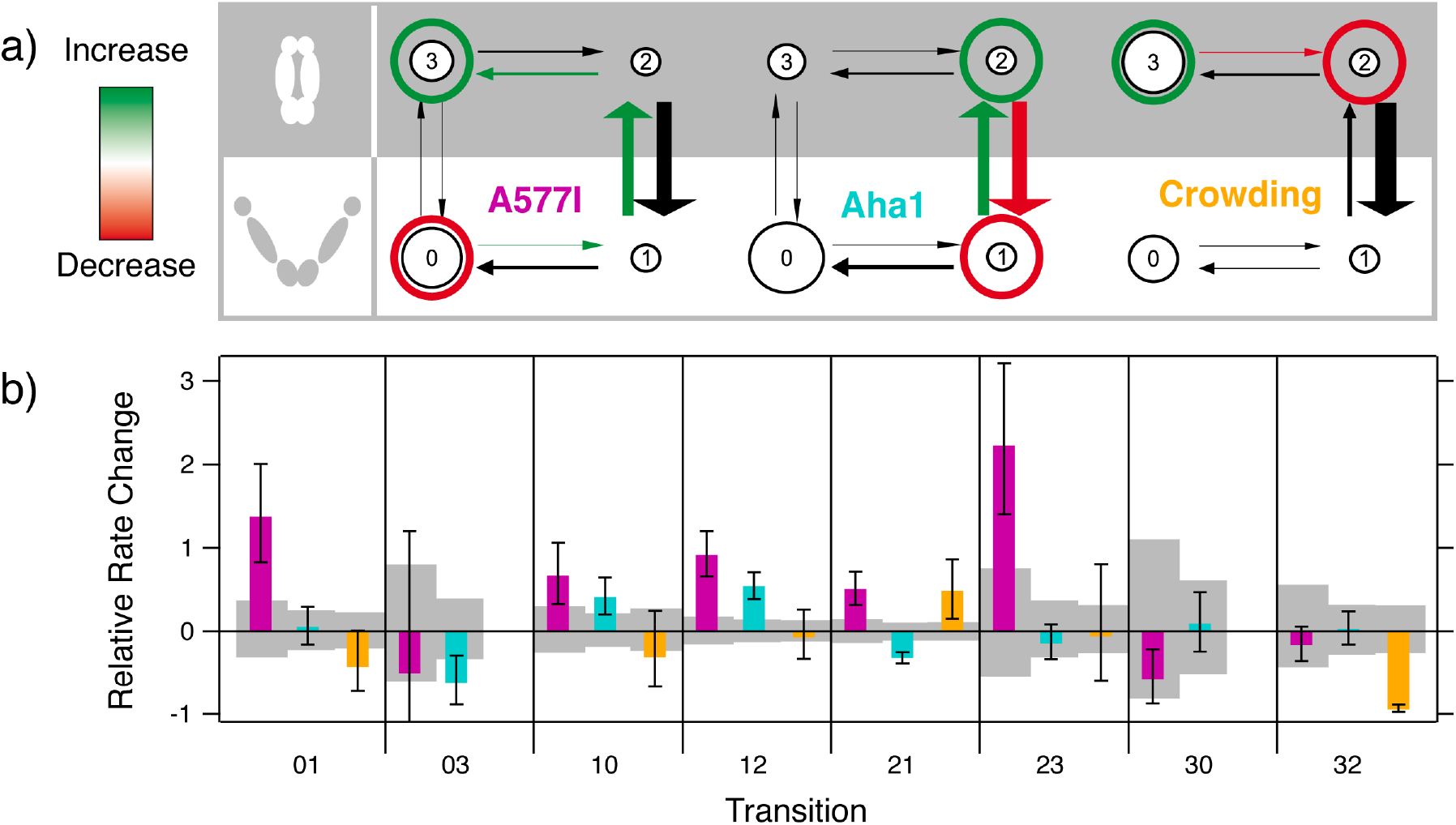
Different conformational kinetics cause similar thermodynamics. (a) Kinetic rate models observed for the point mutant A577I, the cochaperone Aha1, the macro-molecular crowding agent Ficoll00 - each compared to the reference data: wild-type, no Aha1, no crowding, respectively. Significant differences to the reference are highlighted in red and green. Conformational kinetics are described by four states: states 0,1 represent open conformations, and 2,3 are closed conformations. Large and small arrows and circles indicate the size of rates and populations, respectively. For crowding, only 6 links are found, as discussed previously for experiments in the absence of ATP (29,44). (b) The relative rate change under the three conditions in (a) with respect to the reference. Gray boxes show the 95% confidence interval of the reference data. Transition names and color code as in (a). All values are listed in **Table S1.** The molecule counts are the same as for **Fig. 2**.

In the case of cochaperon Aha1, the shift in the FRET efficiency histogram originates from opposing changes of the fast rates between states 1 and 2 (**Fig. S3**). This is in contrast to the effect of the C-terminal point mutation A577I, which collectively accelerates the 3-step pathway out of state 0 to state 3. Note that a kinetic model with only three links - similar to the model for crowding in **Fig. 3a**) **right** - is statistically sufficient (according to likelihood ratio testing detailed in Ref (29) SI point 1.2) to describe the observed kinetics of the A577I homodimer, implying less kinetically heterogeneous fluctuations (**Fig. S4**). This is further evidence in support of an overall stiffened structure of the A577I homodimer with a smoothened (less rough) energy surface, leading to relatively streamlined conformational transitions rather than extensive random walks. Under macro-molecular crowding, changes of the conformational dynamics are visible already from the stationary distributions: as shown in **Fig. 2a**, the low and high FRET populations are most separated in this case. This is indicative of fast fluctuations, at or below the timescale of the sampling rate (5Hz) that are slowed down at higher viscosity. Still, transitions between open and closed conformations are regularly observed in the experiment (**Fig. S5**). For the fully resolved kinetics, the main difference is observed for the rates between the closed states 2 and 3. This agrees with the increase of the closed population under macro-molecular – but not small molecular – crowding. Altogether, **Fig. 3** shows three completely different kinetic effects that underlie seemingly analogous ensemble behavior.

Based on the complete kinetic rate models, we can now deduce the impact on free energies along a specific spatial reaction coordinate, here the N-terminal extension (**Fig. 4**). The point mutation A577I causes an asymmetric destabilization of open conformations. Aha1 leads to simultaneous destabilization of open conformations and stabilization of closed conformations, whereas macro-molecular crowding only stabilizes closed conformations. Importantly, this information is not accessible from the steady-state distributions in **Fig. 2a**.

**Fig. 4:**
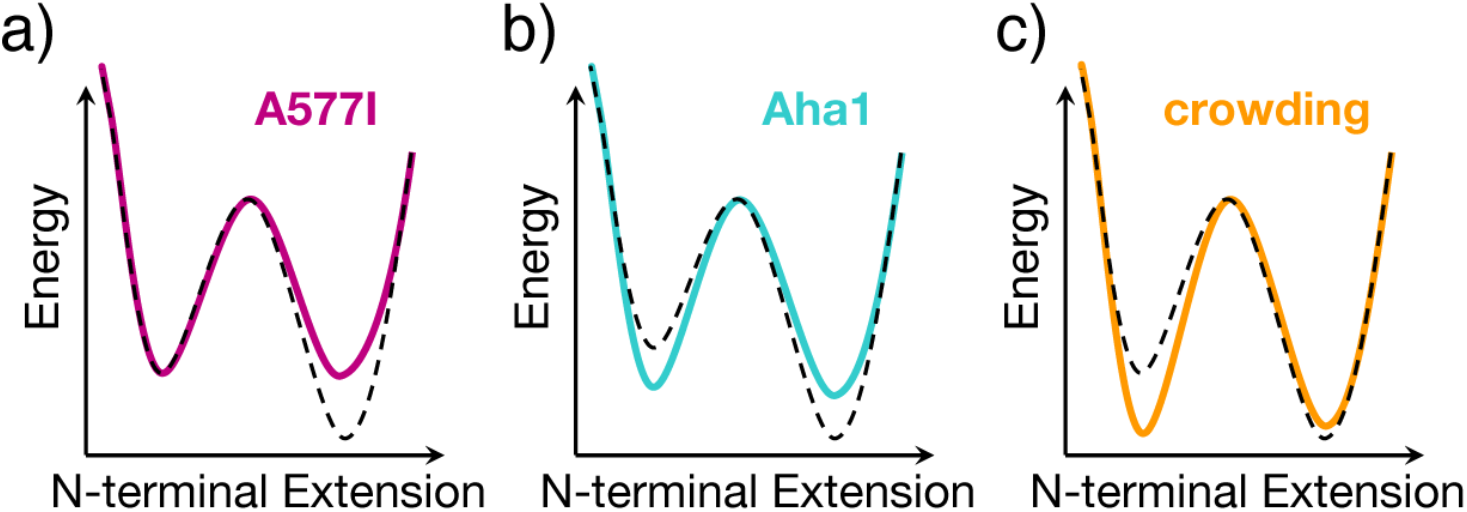
Three contrasting effects on Hsp90’s conformational energy landscape. (a) the open conformation is destabilized by the A577I mutation. (b) Aha1 inversely affects both equilibria. Whereas macro-molecular crowding (c) stabilizes the closed conformation. The dashed black line indicates the reference. This mechanistic information was obtained from all six rate models represented in **Fig. 3**, it is not accessible from **Fig. 2c** alone.

### ATPase stimulation – to varying degrees

The collective shift towards closed conformations is accompanied by an overall increase in ATPase activity under all three conditions (**Fig. 5**): 7-fold for A577I, 17-fold for Aha1, 4-fold for crowding, respectively. Thus, the increase in ATP hydrolysis rate does not reflect the increase in the closed population observed in **Fig. 2c**. This is a first indication of causalities that involve more than just the occurrence of closed conformations. In the following, we dissect the molecular origins of the increased ATPase activity. The effect of macro-molecular crowding can serve as an estimate of the ATPase stimulation caused exclusively by the relative stabilization of closed conformations, because a biochemical interaction of sucrose was excluded in control experiments (see above). Remarkably, the entirely non-specific interaction leads already to a considerable ATPase acceleration of a factor 4. This supports the wide-spread notion that the closed conformation represents Hsp90’s active state (45). But, in comparison to the biochemical effect of the cochaperon Aha1, the stimulation by crowding is still modest, despite the much larger closed population. Specifically, the 3.7-fold increased closed population, comes with a 4-fold increased ATPase activity, whereas in the presence of Aha1 already a 1.6-fold increased closed population is accompanied by a 17-fold ATPase stimulation. The fact that Aha1 induces the smallest increase in closed population, but under the same conditions the largest ATPase stimulation, highlights the functional importance of *specific* contacts between Aha1 and Hsp90, which are responsible for 88% of the ATPase stimulation by Aha1.

**Fig. 5:**
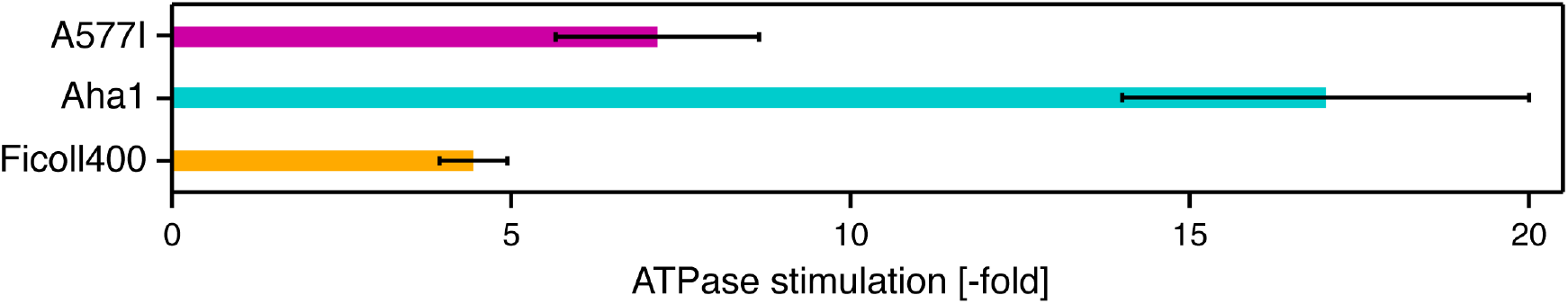
ATPase stimulation by local and global modulations: by the point mutation A577I (34), in the presence of 3μM Aha1 (46), or 20wt% Ficoll400, each normalized by the activity of the wild type, without Aha1, without crowding, respectively.

In the case of the A577I mutant a 2.9-fold increased closed population comes with a 7-fold ATPase amplification. This could result from the mentioned hindrance of extremely open states indicating a conformational stiffening, restricting Hsp90’s native flexibility. In other words, not only changes in the equilibrium, but also changes in the kinetics affect the ATPase activity as discussed below.

## Discussion

Herein we compare three types of Hsp90 modulations, spanning a wide range from a site-specific point mutation, via cochaperon binding, to completely non-specific macro-molecular crowding. All three modulations provoke a similar steady-state behavior, namely an increase in Hsp90’s closed conformation and in Hsp90’s ATPase rate. But significant differences in their kinetics, which could be revealed by single-molecule FRET. This can be rationalized by the emerging picture of yeast Hsp90, as a very flexible dimer that relies critically on external assistance (e.g. by cochaperones) to control this non-productive flexibility. For example, Hsp90’s ATPase function requires the concerted action of the N-terminal nucleotide binding pocket with the ATP lid and distant elements such as the catalytic loop of the middle domain and parts of the opposite N-domain (the N-terminal β1-α1 segment). These elements – also called *the catalytic unit (45)* - however, are very flexible, such as the entire multi-domain dimer. Consequently, anything that constrains this flexibility and confines Hsp90 in a more compact conformation, has a high potential to increase the combined probability for such a concerted action - be it by specific or even non-specific interaction.

**Fig. 6** shows that this can be understood as a direct result of combinatorics: in a flexible protein such as Hsp90, the catalytically active elements have many translational and rotational degrees of freedom. Thus, the probability for a certain hydrolysis-competent conformation is very small. It is however increased dramatically by conditions that constrain these degrees of freedom – even non-specifically – and localize the catalytically active elements. This notion can be further extended to cochaperone binding, and it also implies mutual effects upon client interaction. We conclude, while Hsp90’s flexibility may facilitate its numerous interactions with diverse clients and cochaperones, the flexibility itself has substantial off-state character regarding the ATPase function of Hsp90.

**Fig. 6:**
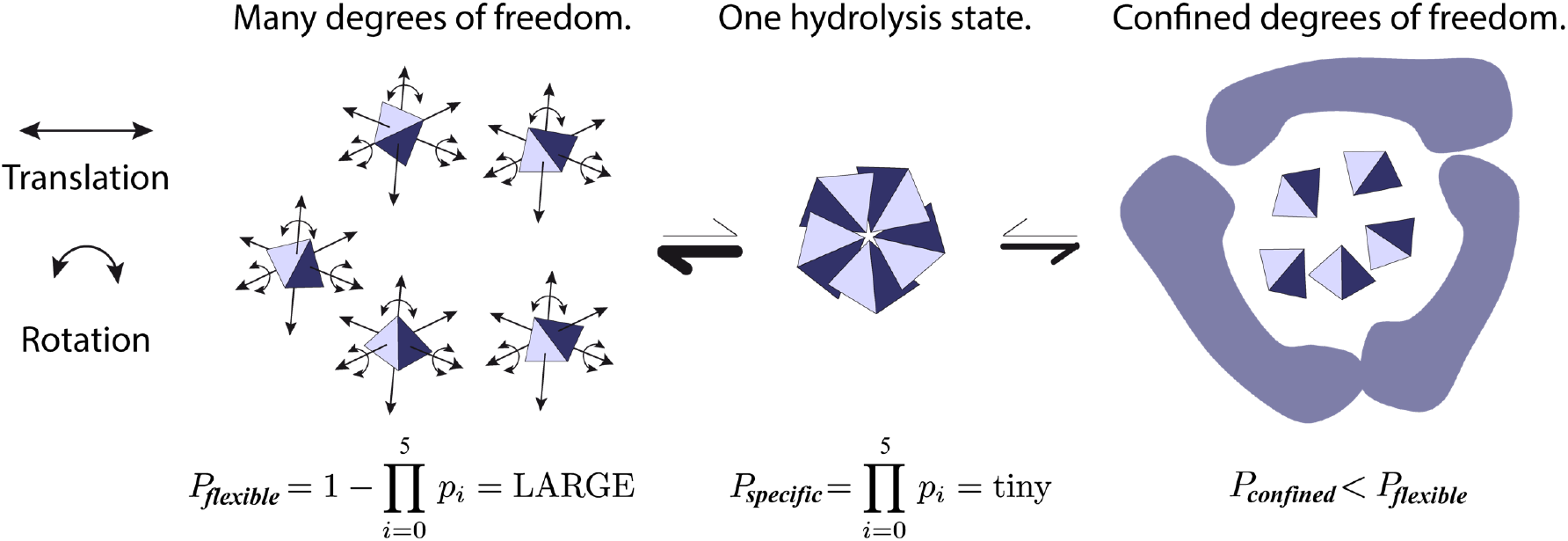
Stimulation of protein function by conformational confinement - a result of combinatorics. In Hsp90 the hydrolysis-competent state consists of multiple flexible elements (illustrated as tetrahedrons) with many degrees of freedom (double-headed arrows), which all have to adopt a specific 3D configuration. The probability *P_specific_* for all elements with *p_i_* to be in this specific configuration is tiny (center), compared to the combined probability for all other accessible configurations (left). Any specific or non-specific confinement (right) decreases the degrees of freedom of the individual elements, thus leading to a relative stabilization of the hydrolysis-competent state. (For clarity, arrows indicating degrees of freedom were omitted on the right.)

A closer look at the regulation of Hsp90’s conformational energy landscape by single-molecule FRET shows the many ways to reach similar ensemble results. The point mutation A557I destabilizes the open conformation, most probably by preventing access to a subset of the conformational ensemble of open states. Macro-molecular crowding stabilizes the closed conformation by simple, sterical confinement. The cochaperone Aha1 combines both mechanisms with additional specific rearrangements. In theory, all three modulations can lead to the exactly same thermodynamic observation, and in fact we observe very similar steady-state distributions. Nevertheless, the kinetics may still vary significantly – as experimentally demonstrated herein. This is an important mathematical fact that holds true for all protein systems. Moreover, our findings indicate that regulation by cochaperones - and protein-protein interactions in general - can have far-reaching thermodynamic, kinetic, and functional consequences. Some of them can possibly be mimicked by individual point mutations, but other consequences might be missed. Lastly, as shown in this work, already non-specific, purely physical macro-molecular crowding has strong effects on thermodynamics, kinetics and function, therefore caution is advised when relating *in vitro* findings to *in vivo* function. As demonstrated herein, in many *in vitro* experiments macro-molecular crowding could easily be included.

In conclusion, we demonstrated herein three ways of protein regulation, ranging from site-specific localized to global modulations. All three show very similar thermodynamic observations, which are, however, caused by clearly different conformational kinetics. This is direct evidence for the importance of kinetics, and of a *dynamic* structure-function relationship in proteins. The reduction of non-productive structural flexibility stimulates Hsp90’s ATPase function – even by entirely non-specific means. Our findings demonstrate that functional stimulation as a result of conformational combinatorics plays an important role in protein regulation. We anticipate that such conformational confinement - by localized *or* global modulations - is an important mechanistic concept with wide-spread implications for protein function in diverse systems.

## Supporting information

Supplement_SchmidHugel

## Methods

### Protein construct preparation

Yeast Hsp90 dimers (UniProtKB: P02829) with a C-terminal coiled-coil motif (kinesin neck region of *D. melanogaster*) were used to avoid dissociation at low concentrations (28). Previously published cysteine positions (41) allowed for specific labeling with donor (61C) or acceptor (385C) fluorophores (see below). Point mutation A577I was introduced using QuikChange Lightning Site-Directed Mutagenesis Kit (Agilent Technologies). The constructs were cloned into a pET28b vector (Novagen, Merck Biosciences, Billerica, MA). They include an N-terminal His-tag followed by a SUMO-domain for later tag cleavage. The QuickChange Lightning kit (Agilent, Santa Clara, CA) was used to insert an Avitag for specific *in vivo* biotinylation at the C-terminus of the acceptor construct. *Escherichia coli* BL21star cells (Invitrogen, Carlsbad, CA) were cotransformed with pET28b and pBirAcm (Avidity Nanomedicines, La Jolla, CA) by electroporation (Peqlab, Erlangen, Germany) and expressed according to Avidity’s *in vivo* biotinylation protocol. The donor construct was expressed in *E. coli* BL21(DE3)cod+ (Stratagene, San Diego, CA) for 3 h at 37°C after induction with 1 mM isopropyl β-D-1-thiogalactopyranoside (IPTG) at OD_600_ = 0.7 in LB_Kana_. A cell disruptor (Constant Systems, Daventry, United Kingdom) was used for lysis in both cases. Proteins were purified as published (46) (Ni-NTA, tag cleavage, anion exchange, size exclusion chromatography). 95% purity was confirmed by SDS-PAGE. Fluorescent labels (Atto550- and Atto647N- maleimide) were purchased from Atto-tec (Siegen, Germany) and coupled to cysteins according to the supplied protocol. If not stated differently, all chemicals were purchased from Sigma Aldrich.

### Single-molecule FRET measurements

smFRET was measured as previously detailed using a home built TIRF setup (29). Hetero-dimers (acceptor + donor) were obtained by 20 min incubation of 1 *μ*M donor and 0.1 *μ*M biotinylated acceptor homodimers in measurement buffer (40 mM Hepes, 150 mM KCl, and 10 mM MgCl_2_, pH7.5) at 47°C. This favors biotinylated heterodimers to bind to the polyethylene glycol (PEG, Rapp Polymere, Tuebingen, Germany) passivated and neutravidin (Thermo Fisher Scientific, Waltham, MA) coated fluid chamber. Residual homodimers are recognized using alternating laser excitation (ALEX) of donor and acceptor dyes (47,48) and excluded from analysis. For optimal interaction affinity with Aha1, measurements were performed in low salt buffer (40mM Hepes, 20mM KCl, 5mM MgCl_2_, pH 7.5 with 3.5*μ*M Aha1 and 2mM ATP). For comparison, data without Aha1 was measured accordingly. Notably, significant binding was previously found for Aha1 with labeled Hsp90-385C at much lower concentration of 0.3*μ*M (49), which is exactly the dissociation constant reported for unlabeled Hsp90 (50). This implies that, although not directly detectable in the experiment, Hsp90 exists predominantly in complex with Aha1 under the used conditions. A577I/wt constructs were created through monomer exchange (see above). They are distinguished from both kinds of homodimers through the fluorescence signal (donor+acceptor). Measurements were performed in measurement buffer plus 2mM ATP if not stated differently. Macro-molecular crowding was mimicked by 20wt% polymeric sucrose, known as Ficoll 400 (Sigma Aldrich) in measurement buffer if not stated differently.

### Activity Assay

ATPase activity was measured at 37°C coupled to NADH oxidation, which was followed as a decrease in absorption at 340nm using an ATP regenerative assay similar to (51): 0.2mM NADH DiNa, Roche; 2mM phosphoenol pyruvate K-salt, Bachem; 2 U/ml pyruvate kinase, Roche; 10 U/ml lactate dehydrogenase, Roche; in 40mM Hepes, 150mM KCl, 10mM MgCl_2_, pH 7.5).

## Acknowledgements

The cochaperone Aha1 was a kind gift of Dr. Markus Jahn. This work has been funded in part by the European Research Council through ERC grant agreement no. 681891 and the Deutsche Forschungsgemeinschaft (DFG, German Research Foundation) – Project-ID 403222702 – SFB 1381. SS acknowledges the Postdoc. Mobility fellowship no. P400PB_180889 by the Swiss National Science Foundation.

