## Supplement_SchmidHugel for "Same Equilibrium. Different Kinetics. Protein Functional Consequences"

### SUPPLEMENTARY INFORMATION – Schmid, Hugel

This file contains Table S1 and Figures S1-S5.

***Table S1: Quantitative values of transition rates and their confidence intervals.*** *All values included in Fig. 3 are specified in Hz, following the nomenclature of the main text, i.e. rate '01' specifies the transition rate constant from state '0' to state '1'. MLE: maximum likelihood estimator of the specified rate constant; +CI / -CI: 95% confidence bound in positive/negative direction, respectively. [*] In the specified one case, -CI is undefined due to asymptotical behavior of the likelihood ratio (i.e. not crossing the chi-squared threshold of 3.81).*

| | Rate | MLE | +CI | -CI |
|---|---|---|---|---|
| **A577I:** | 01 | 0,0381 | 0,0101 | 0,0087 |
| | 03 | 0,0018 | 0,0060 | undef. [*] |
| | 10 | 0,2404 | 0,0569 | 0,0491 |
| | 12 | 0,7949 | 0,1182 | 0,1054 |
| | 21 | 1,0453 | 0,1452 | 0,1311 |
| | 23 | 0,1933 | 0,0592 | 0,0494 |
| | 30 | 0,0075 | 0,0063 | 0,0053 |
| | 32 | 0,0404 | 0,0107 | 0,0093 |
| Reference: | 01 | 0,0160 | 0,0060 | 0,0050 |
| | 03 | 0,0035 | 0,0028 | 0,0021 |
| | 10 | 0,1442 | 0,0440 | 0,0366 |
| | 12 | 0,4147 | 0,0758 | 0,0654 |
| | 21 | 0,6925 | 0,1049 | 0,0942 |
| | 23 | 0,0599 | 0,0456 | 0,0325 |
| | 30 | 0,0176 | 0,0194 | 0,0144 |
| | 32 | 0,0482 | 0,0273 | 0,0208 |
| **Aha1:** | 01 | 0,0218 | 0,0049 | 0,0044 |
| | 03 | 0,0023 | 0,0020 | 0,0016 |
| | 10 | 0,3531 | 0,0585 | 0,0524 |
| | 12 | 0,9828 | 0,1079 | 0,0987 |
| | 21 | 1,2728 | 0,1289 | 0,1193 |
| | 23 | 0,1860 | 0,0492 | 0,0424 |
| | 30 | 0,0111 | 0,0038 | 0,0034 |
| | 32 | 0,0277 | 0,0056 | 0,0051 |
| Reference: | 01 | 0,0207 | 0,0052 | 0,0046 |
| | 03 | 0,0061 | 0,0024 | 0,0020 |
| | 10 | 0,2497 | 0,0543 | 0,0468 |
| | 12 | 0,6376 | 0,0942 | 0,0833 |
| | 21 | 1,8700 | 0,1991 | 0,2000 |
| | 23 | 0,2172 | 0,0813 | 0,0674 |
| | 30 | 0,0101 | 0,0063 | 0,0052 |
| | 32 | 0,0269 | 0,0087 | 0,0074 |
| **crowding:** | 01 | 0,0034 | 0,0026 | 0,0018 |
| | 10 | 0,0176 | 0,0143 | 0,0092 |
| | 12 | 0,0961 | 0,0344 | 0,0269 |
| | 21 | 0,4184 | 0,1071 | 0,0943 |
| | 23 | 0,0442 | 0,0406 | 0,0251 |
| | 32 | 0,0014 | 0,0014 | 0,0008 |
| Reference: | 01 | 0,0059 | 0,0014 | 0,0012 |
| | 10 | 0,0256 | 0,0071 | 0,0060 |
| | 12 | 0,1036 | 0,0140 | 0,0126 |
| | 21 | 0,2818 | 0,0320 | 0,0299 |
| | 23 | 0,0469 | 0,0149 | 0,0124 |
| | 32 | 0,0244 | 0,0076 | 0,0064 |

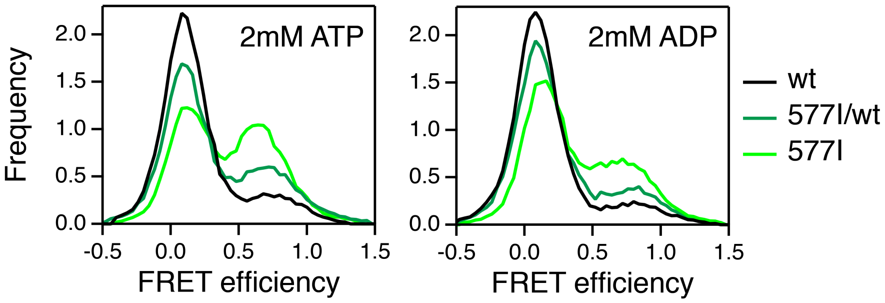


***Figure S1: The additive effect of the A577I mutation on Hsp90's shift towards closed conformations.*** *FRET efficiency histograms for the wild-type (wt), hetero-dimer (577I/wt) and homo-dimer (577I), under ATP and ADP conditions as indicated. Histogram integrals are normalized to unity.*

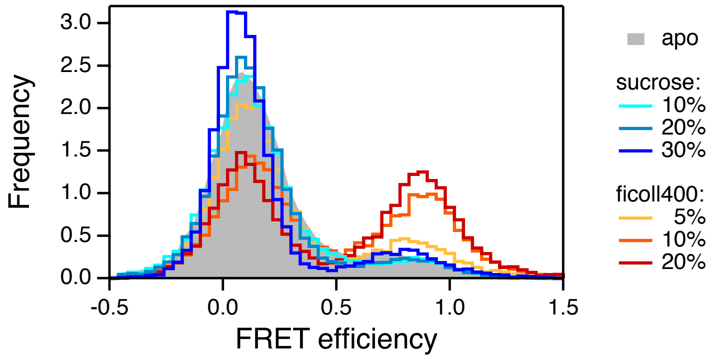


***Figure S2: The differing effect of macro-molecular and small molecular crowding on Hsp90's conformational equilibrium.*** *FRET efficiency histograms in the absence of nucleotides (apo), and with varied weight percent sucrose or Ficoll400, as indicated. Histogram integrals are normalized to unity.*

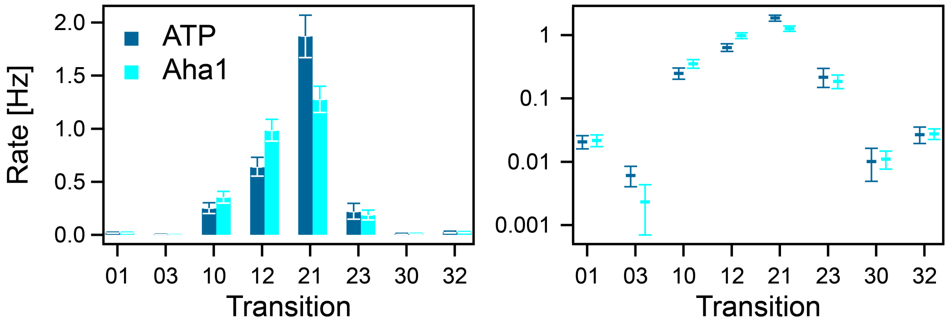


***Figure S3: Rate constants and 95% confidence intervals of Hsp90's opening and closing transitions:*** *in the presence of 2mM ATP (ATP), and in the presence of 2mM ATP + 3.5μM Aha1 (Aha1). Left panels in linear scale, right panels in logarithmic scale.*

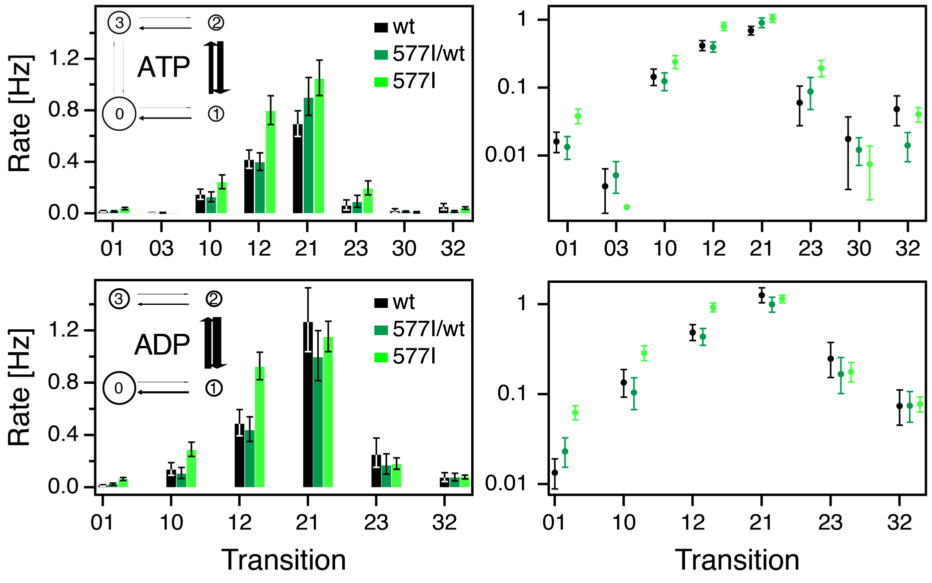


***Figure S4: Rate constants and 95% confidence intervals of Hsp90 opening and closing transitions*** *for the wild-type (wt), and the A577I hetero-dimer (577I/wt), and homo-dimer (577I), under ATP and ADP conditions as indicated. Left panels in linear scale, right panels in logarithmic scale.*

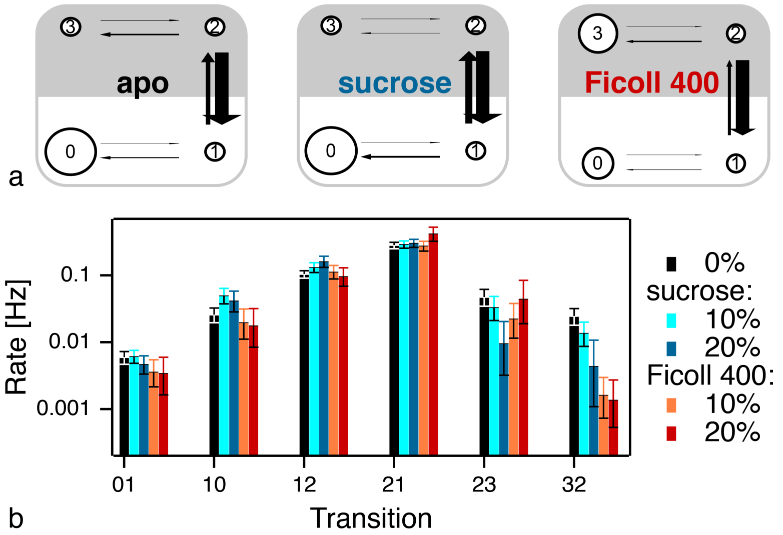


***Figure S5: The effect of crowding on Hsp90's opening and closing transitions: (a)*** *Kinetic state models for wt Hsp90 and 20% sucrose and 20% Ficoll400 (left to right): closed states, gray; open states, white; population represented by circle size; transition rate represented by arrow width.* ***(b)*** *Transition rates with 95% confidence intervals in logarithmic scale.*
